# Validation and Implementation of CLIA-Compliant Whole Genome Sequencing (WGS) in Public Health Laboratory

**DOI:** 10.1101/107003

**Authors:** Varvara K. Kozyreva, Chau-Linda Truong, Alexander L. Greninger, John Crandall, Rituparna Mukhopadhyay, Vishnu Chaturvedi

**Author notes:** Author Contributions: Conceptualization, data analysis and original draft (VKK, VC), data collection and analysis (VKK, CLT, ALG), software code (RM), draft review (VKK, CLT, RM, ALG, VC), project administration and supervision (VC).

## Abstract

**Background:** Public health microbiology laboratories (PHL) are at the cusp of unprecedented improvements in pathogen identification, antibiotic resistance detection, and outbreak investigation by using whole genome sequencing (WGS). However, considerable challenges remain due to the lack of common standards.

**Objectives:** 1) Establish the performance specifications of WGS applications used in PHL to conform with CLIA (Clinical Laboratory Improvements Act) guidelines for laboratory developed tests (LDT), 2) Develop quality assurance (QA) and quality control (QC) measures, 3) Establish reporting language for end users with or without WGS expertise, 4) Create a validation set of microorganisms to be used for future validations of WGS platforms and multi-laboratory comparisons and, 5) Create modular templates for the validation of different sequencing platforms.

**Methods:** MiSeq Sequencer and Illumina chemistry (Illumina, Inc.) were used to generate genomes for 34 bacterial isolates with genome sizes from 1.8 to 4.7 Mb and wide range of GC content (32.1%-66.1%). A customized CLCbio Genomics Workbench - shell script bioinformatics pipeline was used for the data analysis.

**Results:** We developed a validation panel comprising ten *Enterobacteriaceae* isolates, five gram-positive cocci, five gram-negative non-fermenting species, nine *Mycobacterium tuberculosis*, and five miscellaneous bacteria; the set represented typical workflow in the PHL. The accuracy of MiSeq platform for individual base calling was >99.9% with similar results shown for reproducibility/repeatability of genome-wide base calling. The accuracy of phylogenetic analysis was 100%. The specificity and sensitivity inferred from MLST and genotyping tests were 100%. A test report format was developed for the end users with and without WGS knowledge.

**Conclusion:** WGS was validated for routine use in PHL according to CLIA guidelines for LDTs. The validation panel, sequencing analytics, and raw sequences will be available for future multi-laboratory comparisons of WGS in PHL. Additionally, the WGS performance specifications and modular validation template are likely to be adaptable for the validation of other platforms and reagents kits.

## Introduction

Clinical and public health microbiology laboratories are undergoing transformative changes with the adoption of whole genome sequencing (WGS) [1, 2]. For several years, leading laboratories have published proof-of-concept studies on WGS-enabled advances in the identification of pathogens, antibiotic resistance detection, and disease outbreak investigations [3–6]. The technologies also referred to as next generation sequencing (NGS) have yielded more detailed information about the microbial features than was possible using a combination of other laboratory approaches. Further developments of WGS platforms had allowed remarkable in-depth inquiry of pathogenic genomes for the discovery of genetic variants and genome rearrangements that could have been missed using other DNA methods [3, 7, 8]. The enhanced investigations of disease outbreaks have led to new understanding of transmission routes of infectious agents [9–11]. WGS-enabled metagenomics and microbiome discoveries have revealed a new appreciation for the role of microbes in health and disease [12–15]. The innovations are continuing at such an unprecedented pace that WGS is expected to become an alternative to culture-dependent approaches in the clinical and public microbiology laboratories [16–18].

Notwithstanding its promises, several challenges remain for the adoption of WGS in microbiology laboratories [19–22]. The accelerated obsolescence of the sequencing platforms presents several obstacles in bridging the gap between research and routine diagnostics including standardizations efforts [23]. The downstream bioinformatics pipelines are also unique challenges for the microbiology laboratory both in terms of infrastructure and skilled operators [24–27]. Overall, WGS ‘wet bench-dry bench’ workflow represents an integrated process, which is not easily amenable to the traditional quality metrics used by the microbiology laboratories [27–29]. The capital investments and recurring costs of WGS for clinical laboratories although rapidly declining still remain relatively high to allow multi-laboratory comparisons for the standardization of the analytical parameters. Finally, the regulatory agencies have not yet proposed WGS standard guidelines for the clinical microbiology [30], and external proficiency testing programs are still in development for the clinical and public health microbiology laboratories [31, 32].

There are other notable recent developments towards standardization and validation of next generation sequencing in clinical laboratories. The US Centers for Disease Control and Prevention (CDC) sponsored the Next-generation Sequencing: Standardization of Clinical Testing (Nex-StoCT) workgroup to propose quality laboratory practices for the detection of DNA sequence variations associated with heritable human disorders [33, 34]. The workgroup developed principles and guidelines for test validation, quality control, proficiency testing, and reference materials. Although not focused on infectious diseases, these guidelines provide a valuable roadmap for the implementation of WGS in clinical microbiology and public health laboratories. The College of American Pathologists’ (CAP) published eighteen requirements in an accreditation checklist for the next generation sequencing analytic (‘wet bench’) and bioinformatics (‘dry bench’) processes as part of its’ molecular pathology checklist [30]. These ‘foundational’ accreditation requirements were designed to be broadly applicable to the testing of inheritable disorders, molecular oncology, and infectious diseases. Along the same lines, the feasibility of *in silico* proficiency testing has been demonstrated for NGS [35]. Clinical and Laboratory Standards Institute (CLSI) has updated its’ “Nucleic acid sequencing methods in diagnostic laboratory medicine” guidelines with considerations specific to the application of next generation sequencing in microbiology [36]. Thus, a broad technical framework is now available to design WGS validation protocols that will be most relevant for the clinical and public health laboratories. Our aims for the current study were to establish performance metrics for the workflow typical in the microbiological public health laboratories, design modular templates for the validation of different platforms and chemistries, finalize user-friendly report format, and identify a set of bacterial pathogens that could be used for WGS validation and performance assessments.

## Methods

### Bacterial isolates and sequences

A set of 34 bacterial isolates representing typical workflow in the PHL, was used for validation and quality control of WGS. These included ten *Enterobacteriaceae* isolates, five gram-positive cocci bacterial pathogens, five gram-negative non-fermenting bacterial pathogens, nine *Mycobacterium tuberculosis* isolates and five miscellaneous bacterial pathogens (Table 1). This Whole Genome Shotgun project has been deposited at GenBank under the accession MTFS00000000-MTGZ00000000. The version described in this paper is version MTFS01000000-MTGZ01000000. Raw and assembled sequences are available for download (see Supplementary Table 1 for the accession numbers).

**Table 1.** List of strains used for validation and corresponding reference materials.

| Reference materials- NCBI strains |  |  |  |  |  |
| --- | --- | --- | --- | --- | --- |
| MDL ID | Species | Reference |  |  |  |
|  |  | NCBI Strain |  | NCBI Acc# |  |
| C1 | *Escherichia coli* O157:H7 CDC EDL 933 | O157:H7 CDC EDL 933 |  | NZ_CP008957.1 |  |
| C3 | *Escherichia coli* ATCC 8739 | ATCC 8739 |  | NC_010468.1 |  |
| C55 | *Escherichia coli* ATCC 25922 | ATCC 25922 |  | NZ_CP009072.1 |  |
| C4 | *Enterobacter cloacae* ATCC 13047 | ATCC 13047 |  | NC_014121 |  |
| C6 | *Salmonella enterica* ser Typhimurium ATCC 14028 | 14028S |  | NC_016856 |  |
| C5 | *Staphylococcus aureus* ATCC 25923 | ATCC 25923 |  | NZ_CP009361 |  |
| C46 | *Enterococcus faecalis* ATCC 29212 | ATCC 29212 |  | NZ_CP008816 |  |
| C47 | *Staphylococcus epidermidis* ATCC 12228 | ATCC 12228 |  | NC_004461 |  |
| C48 | *Staphylococcus saprophyticus* ATCC 15305 | ATCC 15305 |  | NC_007350 |  |
| C49 | *Streptococcus pneumoniae* ATCC 6305 | ATCC 700669 |  | FM211187 |  |
| C50 | *Pseudomonas aeruginosa* ATCC 27853 | FRD1 |  | NZ_CP010555 |  |
| C51 | *Stenotrophomonas maltophilia* ATCC 13637 | ATCC 13637 |  | NZ_CP008838 |  |
| C52 | *Legionella pneumophila* SG-12 ATCC 43290 | ATCC 43290 |  | NC_016811 |  |
| C53 | *Moraxella catarrhalis* 87A-3084 | ATCC 25240 |  | NZ_CP008804 |  |
| C54 | *Acinetobacter baumannii* ATCC 17945 | AB07 |  | NZ_CP006963 |  |
| C103 | *Bacteroides fragilis* ATCC 25285 | 638R |  | NC_016776 |  |
| C104 | *Haemophilus influenzae* ATCC 10211 | KR494 |  | NC_022356 |  |
| C2 | *Aeromonas hydrophila* ATCC 7966 | ATCC 7966 |  | NC_008570 |  |
| C105 | *Corynebacterium jeikeium* ATCC 43734 | ATCC 43734 |  | GG700813:GG700833 |  |
| C106 | *Neisseria gonorrhoeae* ATCC 49226 | MS11 |  | NC_022240 |  |
| C56 | *Mycobacterium tuberculosis* | H37Rv |  | NC_000962.3 |  |
| C57 | *Mycobacterium tuberculosis* | H37Rv |  | NC_000962.3 |  |
| C58 | *Mycobacterium tuberculosis* | H37Rv |  | NC_000962.3 |  |
| C59 | *Mycobacterium tuberculosis* | H37Rv |  | NC_000962.3 |  |
| C61 | *Mycobacterium tuberculosis* | H37Rv |  | NC_000962.3 |  |
| C65 | *Mycobacterium tuberculosis* | H37Rv |  | NC_000962.3 |  |
| C67 | *Mycobacterium tuberculosis* | H37Rv |  | NC_000962.3 |  |
| C68 | *Mycobacterium tuberculosis* | H37Rv |  | NC_000962.3 |  |
| C69 | *Mycobacterium tuberculosis* | H37Rv |  | NC_000962.3 |  |
| Reference materials- strains sequenced at CDC |  |  |  |  |  |
| MDL ID | Species | Reference raw reads generated by CDC |  | Reference used for mapping |  |
|  |  | CDC Strain | Accession # | NCBI Strain | NCBI Acc# |
| C72 | *Escherichia coli* O121:H19 | 2014C-3857 | SRR1610033 | 2011C-3493 | NC_018658 |
| C73 | *Salmonella enterica* ser Enteritidis | CDC_2010K-1543 | SRR518749 | P125109 | NC_011294.1 |
| C74 | *Salmonella enterica* ser Infantis | 2014K-0434 | SRR1616809 | 1326/28 | NZ_LN649235 |
| C75 | *Salmonella enterica* ser Adelaide | 2014K-0941 | SRR1686419 | P125109 | NC_011294.1 |
| C76 | *Salmonella enterica* ser Worthington | 2012K-1219 | SRR1614868 | P125109 | NC_011294.1 |
| C77* | *Salmonella enterica* ser Saintpaul | 2014K-0875 | SRR1640105 | 14028S | NC_016856 |

**Footnotes:** Green color designates cases when the genome of the strain sequenced by the PHL is available from the NCBI database. Yellow color designates cases when the genome of the strain sequenced by the PHL is NOT available from the NCBI database and an alternative reference genome was used for mapping.
\*P.S.: Sample C77 was sequenced by PHL only for genotyping test accuracy validation. No replicates were done.

### Reference whole genomes

The genome sequences of ATCC strains, isolates characterized by CDC, and other representative isolates were downloaded from NCBI database (http://www.ncbi.nlm.nih.gov/genome/) to be used as reference per the recommendations in the CLSI guidelines [36], (Table 1).

### WGS wet bench workflow

The whole genome sequencing was performed on Illumina MiSeq sequencer (Figure 1). The Nextera XT library preparation procedure and 2x300 cycle MiSeq sequencing kits were used (Illumina Inc., San Diego, CA, USA). Illumina Nextera XT indexes were used for barcoding. Bacterial DNA was extracted using Wizard Genomic DNA Kit (Promega, Madison, WI, USA). The bacterial DNA concentrations were measured using Qubit fluorometric quantitation with Qubit dsDNA BR Assay Kit (Thermo Fisher Scientific, Waltham, MA, USA). The DNA purity was estimated using NanoDrop 2000 UV-Vis Spectrophotometer (NanoDrop Products, Wilmington, DE, USA). The Mastercycler nexus was used for tagmentation incubation and PCR (Eppendorf North America, Hauppauge, NY, USA). The library concentration was measured using Qubit HS kit. DNA library size distribution was estimated using 2100 BioAnalyzer Instrument and High Sensitivity DNA analysis kit (Agilent Technologies, Santa Clara, CA, USA). Ampure beads were used for size selection. Manual normalization of libraries was performed. The PhiX Control V3 sequencing control was used in every sequencing run (Illumina, Inc. San Diego, CA, USA). Genomes were generated with the depth coverage in the range of 15.71x-216.4x (average 79.72x, median 71.55x).

**Figure 1.**
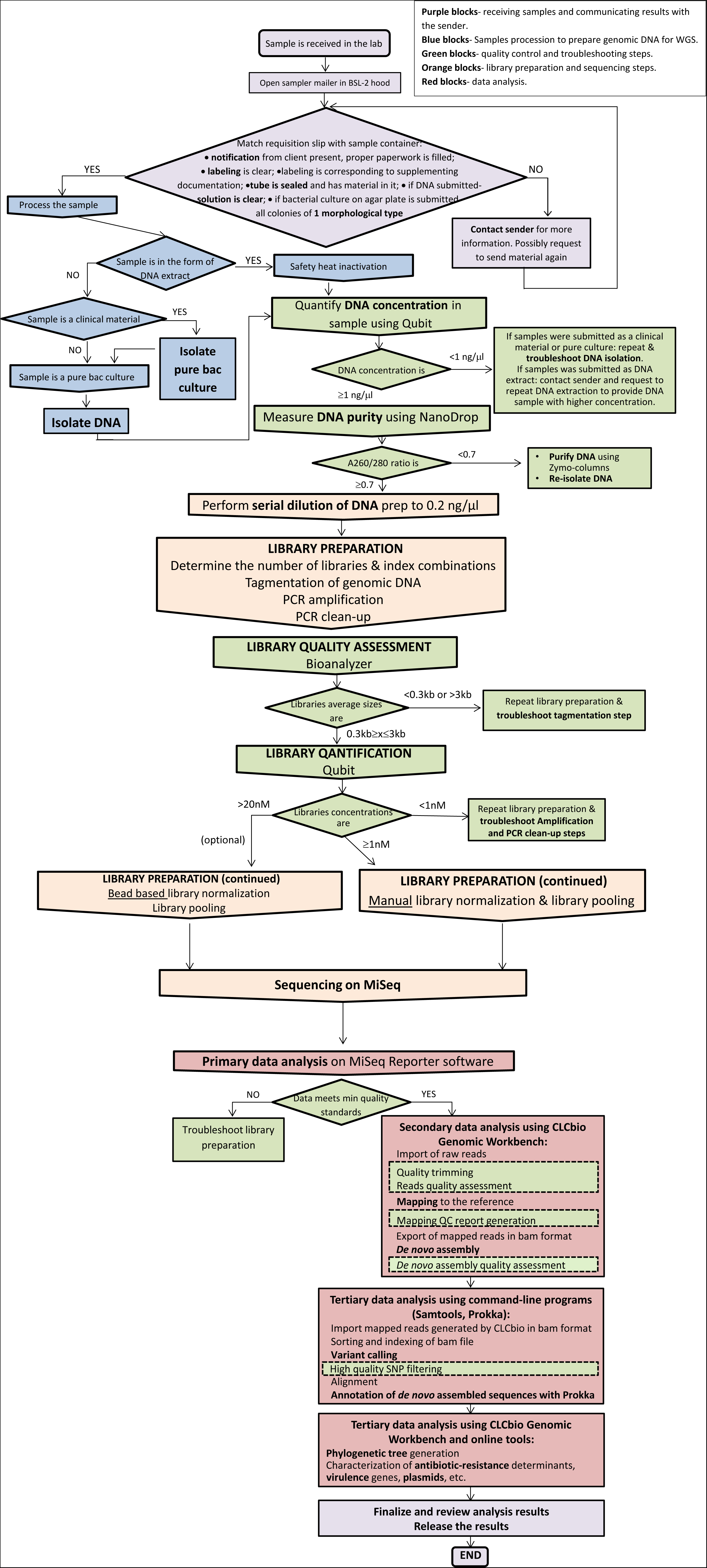
WGS wet and dry bench workflow.

### Bioinformatics pipeline

Paired-end reads were quality trimmed with the threshold of Q30, and then used for mapping to the reference and *de novo* assembly on CLCbio Genomic Workbench 8.0.2 (Qiagen, Aarhus, Denmark). The BAM files generated after mapping to the reference genome were taken through series of software suites to generate the phylogenetic tree. A customized shell script was created to automate the subsequent steps after mapping that included: 1) SNP calling in coding and non-coding genome areas using SAMtools mpileup (v.1.2; [37]); 2) Converting into VCF matrix using bcftools (v0.1.19; http://samtools.github.io/bcftools/); 3) Variants parsing using vcftools (v.0.1.12b; [38]) to include only high-quality SNPs (hqSNPs) with coverage ≥30x, minimum quality > 200; with InDels and the heterozygote calls excluded. 4) Converting SNP matrix into FASTA alignment file for the export back to the CLCbio GW 8.0.2 for the generation of the phylogenetic tree.

<u>hqSNP-based genotyping</u> - The Maximum Likelihood phylogenetic trees were generated based on high-quality single nucleotide polymorphisms (hqSNPs) under the Jukes-Cantor nucleotide substitution model; with bootstrapping.

<u>16S rRNA gene-based identification</u> - Genomes were annotated with prokka v1.1 tool [39] and species identification was performed by comparing 16S rRNA gene sequences against the Ribosomal Database Project (RDP) database [40].

<u>*In silico* MLST</u> - *In silico* multi-locus sequence typing (MLST) was performed using the Center for Genomic Epidemiology (CGE) online tool [41].

<u>ABR genes detection</u> was performed using the CGE ResFinder online resource [42]. ATCC reference strains designated for use as antibiotic susceptibility controls were analyzed for the presence of antibiotic resistance genes. Negative controls were chosen among strains which were described by the CLSI M100-S25 document [43] as susceptible, with no known antibiotic resistance genes. Positive controls were chosen among strains, which according to the CLSI M100-S25 resistance determinants.

### Validation Plan

Thirty-four bacterial isolates were sequenced in triplicate. For between-run reproducibility assessments, all replicates were generated starting from fresh cultures except for *M. tuberculosis* where DNA samples were used. Between run replicates were processed on separate days by different operators. For within run replicates, one DNA extract was used, but independent library preparations were done, with final samples being included in one sequencing run.

## Results

A number of CLIA-required quality parameters were adopted with some modification for validation on WGS (Table 2). The modular validation template and a summary of performed here WGS validation for 34 bacterial isolates are presented in figure 2.

**Table 2.**
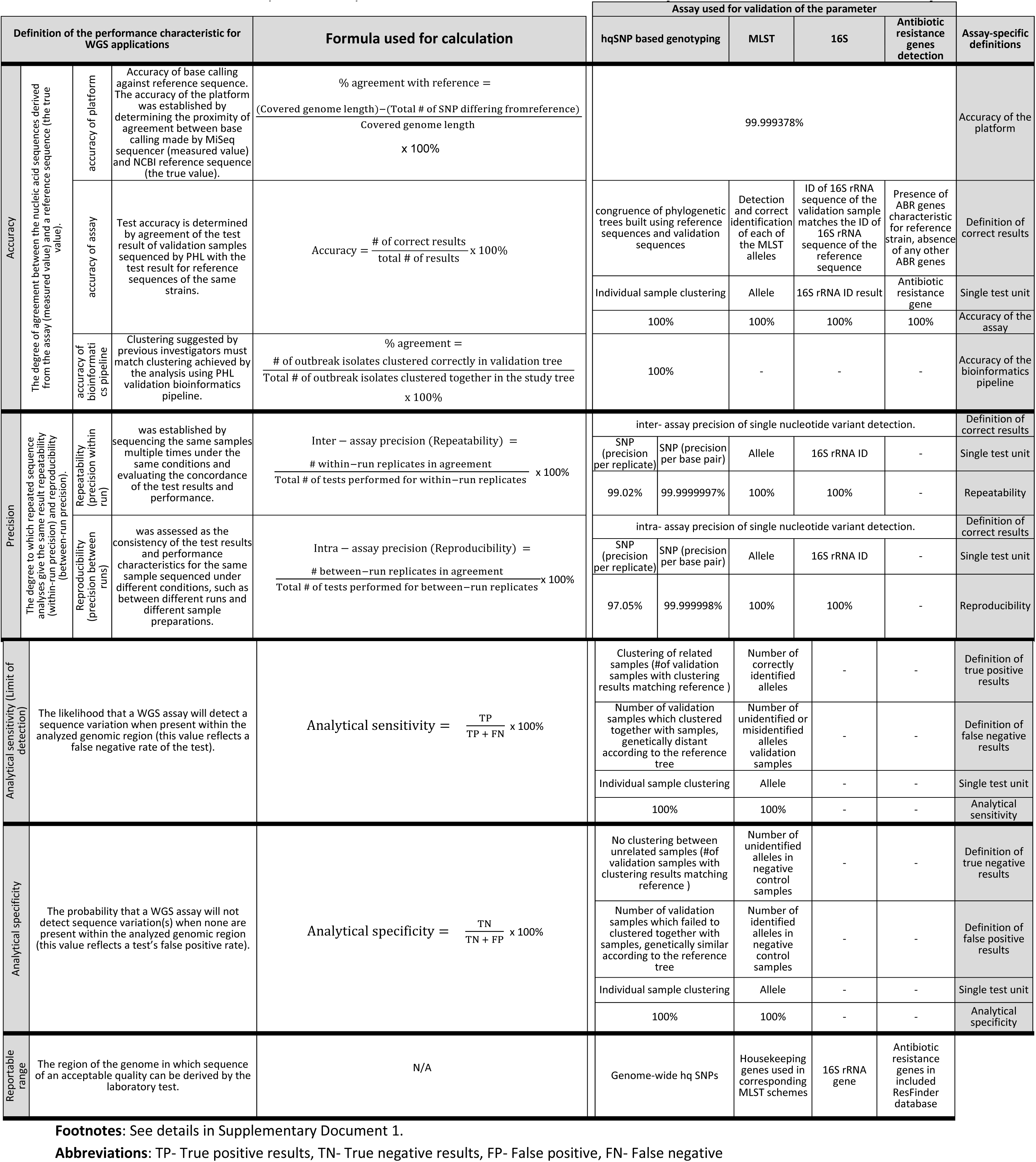
Performance characteristics, definitions, and formulas used in validation. Summary of the validation for different assays.

**Figure 2.**
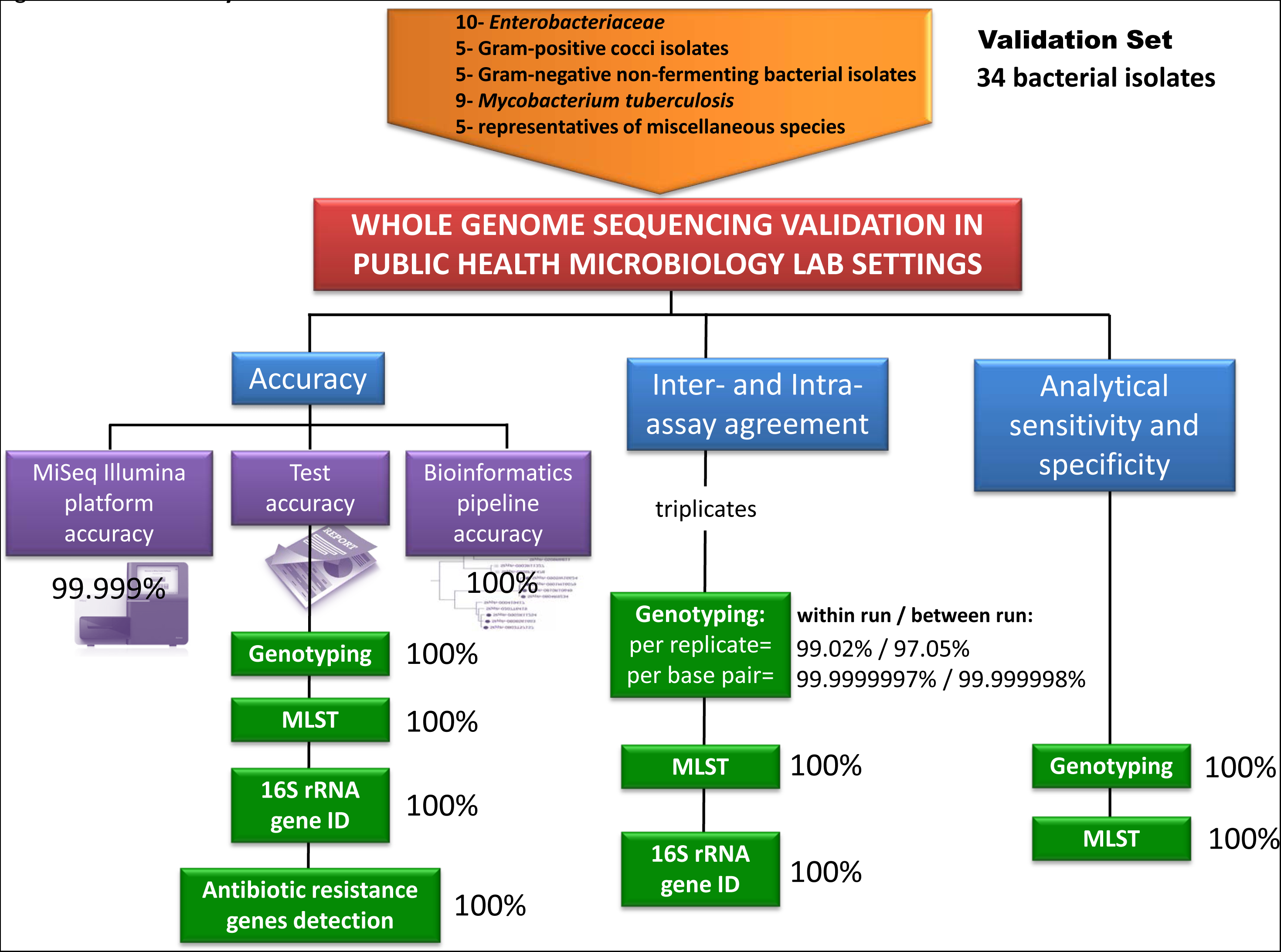
The summary of the WGS validation.

### Accuracy of WGS

The accuracy of WGS was divided into three components: platform accuracy, assay accuracy, and bioinformatics pipeline accuracy.

<u>Platform accuracy</u> - Platform accuracy was assessed as the accuracy of identification of individual base pairs in the bacterial genome. The accuracy of the platform was established by determining the proximity of agreement between base calling made by MiSeq sequencer (measured value) and NCBI reference sequence (the true value). We determined MiSeq Illumina platform accuracy by mapping generated reads to the corresponding reference sequence and identifying Single Nucleotide Polymorphisms (SNPs). Few validation samples differed from reference genome by several SNPs.

However, 99% (324 out of 327) of those SNPs were reproducible among all five replicates we have sequenced for each sample. Since sequencing errors are random between different library preparations and it is unlikely that the same erroneous SNP will occur in all 5 replicates, we can conclude that those discrepancies were not caused by sequencing errors, but most likely were a result of accumulation of mutations in the reference strains or previous sequencing mistakes in the reference sequence. In both cases, whether we take into the account all SNPs detected between validation and reference sequence, or only those SNPs which don’t appear in all of the replicates (true sequencing errors), we observed > 99.999% agreement of generated whole genome sequences with the reference sequences for each tested sample.

<u>Assay accuracy</u> - Assay accuracy was determined by an agreement of the assay result for the validation samples with the assay result for reference sequences of the same strains. Four applications of WGS were used to validate the accuracy of the assay: *in silico* <u>M</u>ultilocus <u>S</u>equence <u>T</u>yping (MLST) assay, 16S rRNA gene species identification (ID) assay, an assay for detection of antibiotic resistance (ABR) genes, and genotyping assay using high-quality <u>S</u>ingle <u>N</u>ucleotide <u>P</u>olymorphisms (hqSNPs).

The definition of the correct result for MLST corresponds to a correct identification of each of the MLST alleles in the validation sequence. For all validation samples each of the sequences of the seven housekeeping genes used in the typing scheme (or 6 genes-for *Aeromonas hydrophilia*) were identified correctly, resulting in 100% allele identification accuracy.

For ABR genes detection the comparison of validation sequences was performed against each entry in the ResFinder database, which at the moment of validation contained sequences of 1719 antibiotic resistance genes, resulting in a total of 1719 tests performed for each validation sample. In negative control samples, all 1719 tests gave negative results. In positive controls, 1 out of 1719 tests gave a positive result, and the rest must remain negative, as expected. Thus, the accuracy of the assay for ABR genes detection was 100%.

For 16S rRNA ID assay, variations only in one gene were detected, so the species ID results as a whole (e.g. “*Escherichia coli*”) was considered as a single test. The identity of 16S rRNA sequence extracted from validation sample showed 100% match with 16S rRNA sequence extracted from the reference sequence.

To assess the accuracy of the genotyping test, phylogenetic trees were built using reference sequences and validation sequences, and resulting trees were compared. For better comparison, we used at least five strains of the same species in the phylogenetic tree. The accuracy of the genotyping test was determined using two approaches: 1) Topological similarity between reference tree and validation tree using Compare2Trees software, and 2) Comparison of clustering pattern of validation tree and reference tree. The phylogenetic trees were generated for five bacterial isolates. All five validation trees had matching clustering patterns and 100% of topological similarity with corresponding reference trees (Supplementary Table 2).

<u>Bioinformatics pipeline accuracy</u> - Accuracy of the bioinformatics pipeline used for hqSNP genotyping was assessed by performing phylogenetic analysis on raw WGS reads of bacterial isolates from well-characterized outbreaks and comparing validation results to the previously published phylogenetic results (Table 3). Two studies, presenting a phylogenetic analysis of outbreaks, caused by the gram-positive pathogen in one study [44] and gram-negative in another study [45] (at least six isolates/study), were used for validation of the bioinformatics pipeline (Figure 3). The clustering of validation tree completely replicated clustering of Study 1 [44] tree (Figure 3A-C), e.g. isolates 4 and 5 were identical and clustered together according to the Study 1, and the same results were shown in validation tree, with isolates 4 and 5 sharing the same node. All conclusions in regards to the genetic relatedness of the isolates that can be drawn from Study 1 tree can also be made from analysis of validation tree 1. The group of related isolates from Study 1 was compared with epidemiologically unrelated isolates suggested by the same study (no tree available from publication). The phylogenetic analysis using the PHL bioinformatics pipeline showed that epidemiologically unrelated isolates did not cluster with the group of outbreak isolates and appeared to be genetically distant (Figure 3D). Thus, the resulting phylogenetic tree produced by our bioinformatics pipeline showed complete concordance with the epidemiological data.

**Table 3.** Summary of the studies used for validation of bioinformatics pipeline.

| **Study** | **Study 1.**<br>SR Harris et al. Lancet Infect Dis 2013;<br>13: 130–36 [44] | **Study 2.**<br>P Leekitcharoenphon et al. PLoS ONE<br>2014; 9(2): e87991 [45] |
| --- | --- | --- |
| **Microorganism** | Methicillin-resistant *Staphylococcus aureus* | *Salmonella enterica* serovar Typhimurium |
| **Source of isolates** | Human | Human |
| **Number of isolates analyzed** | 7 outbreak isolates (1 outbreak cluster) +<br>2 epidemiologically unrelated isolates | 9 outbreak isolates (4 outbreak clusters)<br>+ 2 epidemiologically unrelated isolates |
| **Type of outbreak** | Hospital-associated outbreak | Foodborne outbreaks |
| **ID of the samples in the study which were used for validation** | P1, P2, P3, P4, P16, P21, P25, Identified by Infectious Control Investigation non-outbreak ST1, MRSA identified by searching microbiology database non-outbreak ST772 | 0803T57157, 0808S61603, 0808F31478, 0903R11327, 0811R10987, 0804R9234, 0810R10649, 0901M16079, 0110T17035, 1005R12913, 1006R12965 |
| **Accession ## of corresponding samples** | ERR070045, ERR070042, ERR070043, ERR070044, ERR124429, ERR124433, ERR128708, ERR070041, ERR072248 | ERR277220, ERR277226, ERR277223, ERR277222, ERR277224, ERR277221, ERR277227, ERR277228, ERR277203, ERR277233, ERR277234 |
| **# of clusters in study tree** | 1 | 4 |
| **# of clusters in validation tree** | 1 | 4 |
| **# of outbreak isolates in each cluster in the study tree** | Cluster 1= 7 | Cluster 1= 2, Cluster 2= 3, Cluster 3= 2, Cluster 4= 2 |
| **# of outbreak isolates in each cluster in validation tree** | Cluster 1= 7 | Cluster 1= 2, Cluster 2= 3, Cluster 3= 2, Cluster 4= 2 |
| **# of epidemiologically unrelated isolates in the set** | 2 | 2 |
| **# of epidemiologically unrelated isolates clustered with outbreak isolates** | 0 | 0 |
| **% agreement= (# of outbreak isolates clustered correctly in validation tree)x100%/ (Total # of outbreak isolates clustered together in the study tree)** | $(7 \times 100 / 7) = 100\%$ | $(9 \times 100 / 9) = 100\%$ |

**Figure 3.**
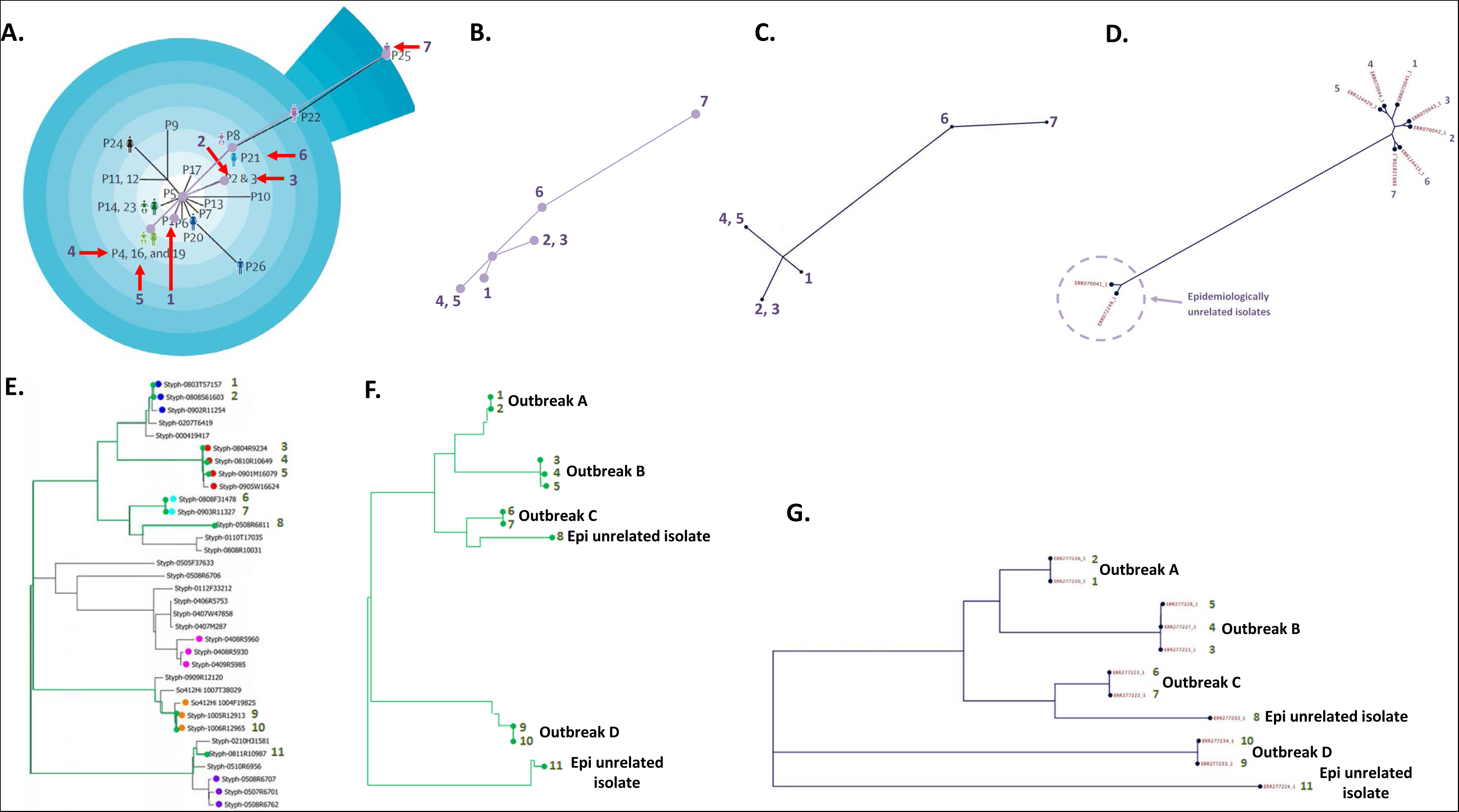
Bioinformatics pipeline validation with two groups of outbreak isolates. **A.** “Study 1 tree”, a phylogenetic tree of outbreak isolates, which was published in the study 1. The isolates from the study which were picked for validation have arrows pointing at them and numbers assigned for purposes of validation (1-7). **B.** A tree representing phylogenetic connections between chosen isolates from original study tree. **C.** “Validation tree 1”, a phylogenetic tree generated using the PHL bioinformatics pipeline. The same isolates in the original tree and validation tree are marked with the same numbers. **D.** Comparison of the group of related isolates (1–7) from Study 1 with epidemiologically unrelated isolates from the same study using the PHL bioinformatics pipeline. **E.** "Study two tree", a phylogenetic tree combining epidemiologically related and nonrelated isolates published in the study 2. The isolates from the study two which were picked for validation marked with green node circles and had numbers 1-11 assigned for purposes of validation. **F.** A tree representing phylogenetic connections between chosen isolates from original study tree. **G.**“Validation tree 2”, a phylogenetic tree generated using the PHL bioinformatics pipeline. The same isolates in the tree from Study 2 and the validation tree are marked with the same numbers.

From the Study 2 [45], we have selected nine isolates, which were representative of 4 independent outbreaks and two isolates were epidemiologically unrelated controls (Figure 3E-G). The clustering of validation tree was identical to the clustering of Study 2 tree. For example, isolates 6 and 7 were a part of the same outbreak, while isolate 8 is an epidemiologically unrelated control used in the study. By epidemiological data and Study 2 tree, the validation tree showed that isolates 6 and seven do cluster together, but not with isolate 8. All observations about the genetic relatedness of the isolates drawn from Study 2 tree could be replicated from the analysis of validation tree 2. In summary, based on analysis of simulated data from both studies accuracy of the pipeline for phylogenetic analysis was 100%.

### WGS repeatability and reproducibility

Repeatability (precision within run) was established by sequencing the same samples multiple times under the same conditions and evaluating the concordance of the assay results and performance. Reproducibility (precision between runs) was assessed as the consistency of the assay results and performance characteristics for the same sample sequenced on different occasions. Thirty-four validation samples each were sequenced three times in the same sequencing run (for repeatability) and in 3 times in different runs (for reproducibility). Between run replicates were processed on different days, altering two operators, as recommended CLSI MM11A document [46]. For within run replicates, one DNA extract was used, but independent library preparations were done, with final samples being included in one sequencing run. Therefore, for each sample, the number of intra-assay replicates and inter-assay replicates were three each, and the total numbers of repeated results were five. All quality parameters [depth of coverage, uniformity of coverage, and accuracy of base calling (Q score)] remained relatively constant within and between runs.

Two methods of evaluating precision were used: evaluation of absolute inter- and intra-assay precision per replicate and evaluation of precision relative to the genome size. One out of 3 within-run replicates of isolate C50 *Pseudomonas aeruginosa* ATCC 27853 had a 1 SNP difference from other within-run replicates (see Supplementary Table 3). All validation samples except C50 yielded identical whole genome sequences for all three within-run replicates. The inter-assay precision was 99.02% as per replicate. Three validation samples had one of the between-run replicates each differing from other between-run replicates. Sample C47 *Staphylococcus epidermidis* ATCC 12228 had one between-run replicate with 2 SNPs difference from other replicates. Samples C49 *Streptococcus pneumoniae* ATCC 6305 and C55 *Escherichia coli* ATCC 25922 each had one of the between-run replicates differing from other replicated sequences by 1 SNP. Intra-assay precision per replicate was 97.05%. If precision per base pair is estimated (in relation to the covered genome size), both inter- and intra-assay precision were > 99.9999%.

We also estimated reproducibility and repeatability for MLST and 16S rRNA ID assays. For MLST total number of alleles analyzed for either within- or between-run replicates was 441. Each single allele in all validation samples was identified consistently among within- and between-run replicates. Within- and between-run replicates had repeatable/reproducible sequences of 16S rRNA gene and resulted in repeatable/reproducible species identification. Within and between run precisions of allele detection and species identification for corresponding assays were 100%.

### WGS Sensitivity and Specificity

Analytical sensitivity and specificity of WGS were estimated for genotyping and MLST.

<u>Genotyping sensitivity and specificity</u> - to estimate analytical sensitivity and specificity of WGS-based genotyping, the hqSNPs phylogenetic trees generated from the validation sequences were compared to the trees generated from the reference sequences for the same strain. All generated validation trees repeated clustering and had 100% of topological similarity with corresponding reference trees, indicating absence false negative or false-positive results in the genotyping test. Both analytical sensitivity and analytical specificity of the hqSNP-based genotyping assay were 100%.

<u>MLST sensitivity and specificity</u> - As described above, using organism-specific MLST databases sequence type of validation sequences and their reference sequences was determined. For MLST number of the true positive results corresponds to the number of alleles correctly identified in the validation samples. For the true negative results, we performed a comparison of validation sequences against MLST databases for unmatched species, e.g. search of alleles for C1 *Escherichia coli* validation sample against MLST database for *Salmonella enterica*. In the latter case, the MLST assay is not supposed to be able to identify any alleles. All alleles in positive validation samples were identified correctly. None of the alleles in negative controls were identified. Both analytical sensitivity and analytical specificity of *in silico* MLST test were 100%.

### WGS reportable range

The following information about the sequenced genome was collected for the reportable range: genome-wide hq SNPs, housekeeping genes used in MLST schemes, 16S rRNA gene, and antibiotic resistance genes included into ResFinder database.

Reporting language was developed to assist interpretation of the results by an end user with or without specific WGS knowledge-the template and examples are provided in the Supplementary Document 1.

### Quality assurance and quality control of WGS

The quality assurance (QA) and quality control (QC) measures were developed as the results of valuation to ensure high quality and consistency of further routine testing using MiSeq Illumina platform. QC must be performed during both pre-analytical (DNA isolation, library preparation), analytical (quality metrics of sequencing run) and post-analytical (data analysis) steps of the WGS. On the stage of data analysis, QC includes three steps: raw read QC, mapping quality QC (or/and *de novo* assembly QC), variant calling QC. PHL should use the WGS validation to establish the thresholds of quality parameters, which can be used in following routine testing to filter out poor quality samples and data and this way minimize the chance of false results. We suggest spiked-in positive and negative controls for routine testing as well as more comprehensive monthly positive and negative controls. Since traditional CLIA rules require the positive and negative control to pass through all the pre-analytical steps, including DNA isolation, laboratory may choose to follow this guidance and perform DNA isolation and sequencing of positive and negative control in each run, or alternatively, implement Individualized Quality Control Plan (IQCP) [as per 42CFR493.1250] and use more economical spiked-in control instead. Type and complexity of positive and negative controls should be determined by each laboratory individually based on specifics of their workflow (most probable source of contamination), type of microorganisms and assays which are most commonly used. Regular and monthly QC practices are summarized in Supplementary Figure 1. The complete QA&QC manual established for WGS applications used in microbiological PHL can be found in Supplementary Document 1.

### Validation Summary

WGS assay was shown to have >99.9% accuracy, >99.9% reproducibility/repeatability, and 100% specificity and sensitivity, which meets CLIA requirements for laboratory-developed tests (LDTs).

## Discussion

This study established the workflow and reference materials for the validation of WGS for routine use in PHL according to CLIA guidelines for LDTs. The validation panel, sequencing analytics, and raw sequences generated during this study could serve as a resource for the future multi-laboratory comparisons of WGS. Additionally, the WGS performance specifications and modular validation template developed in the study could be easily adapted for the validation of other platforms and reagents kits. These results strengthen the concept of unified laboratory standards for WGS enunciated by some professional organizations, including the Global Microbial Identifier (GMI) initiative [30, 31, 33, 47]. A few other groups have also highlighted the challenges and solutions for the implementation of WGS in clinical and public health microbiology laboratories [21, 48].

Using a combination of reference strains and corresponding publicly available genomes, we devised a framework of ‘best practices’ for the quality management of the integrated ‘wet lab’ and ‘dry lab’ WGS workflow (‘pipeline’). The importance of reference materials for validation and QC of wet–and dry-lab WGS processes has been noted earlier [28, 31, 33]. Unlike in human genomics [49], there is no well-established source of reference materials for WGS validation in microbiological PHL. The main challenge of creating customized validation set is the lack of reference materials, in other words, strains that can be easily acquired by the PHLs and which have high-quality well-characterized reference genomes available. While using complete genomic sequences of ATCC strains from NCBI is an option, it is far from being perfect. The genome sequences available from public databases are generated by using different methods, chemistries, platforms, which may yield different error rates, therefore deposited sequences are not guaranteed to be free of such errors. With the perpetual development of new sequencing technologies and improvements in the quality of sequences, it is not unlikely that the genomes sequenced with old methods may appear less accurate than the validation sequences generated by the laboratory during validation. In addition to this, there is a possibility of mutations accumulation in the control strains, e.g. ATCC cultures, which are propagated by the different laboratories. In this sense, there is no gold standard available for use as a reference material for bacterial WGS validation. Nevertheless, NCBI, ENA, and similar public genome depositories remain to be the best resource for the genomic sequences of control strains, which can be used for validation. In future, it would be optimal to have a network/agency/bank which could distribute panels of thoroughly sequenced isolates, with curated and updated genomic sequences available online for WGS validation. In the absence of such resource, we developed a validation set of microorganisms, which can be used for future validations of WGS platforms. Bacterial genomes vary differently in size, GC content, abundance of repetitive regions, and other properties, which affect the WGS results. We created a validation set which reflects the diversity of the microorganisms with various genome sizes and GC-content, which are routinely sequenced by the PHL. Different species of gram-positive and gram-negative microorganisms and *M. tuberculosis* were included to account for the differences in DNA isolation procedures as well.

Samples were validated based on four core elements also reflected in the assay report: 16S rRNA-based species identity, *in silico* MLST, hqSNP phylogenetic analysis, and the presence of AR determinants. Overall, we achieved high accuracy, precision, sensitivity and specificity for all test analytes ranging from 99-100%, which well exceeds 90% threshold for these performance parameters for LDT as per CLIA. These findings are in agreement with recent reports of 93%-100% accuracy in WGS identification, subtyping, and antimicrobial resistance genes detection in a number of pathogens [50–53].

The successful CLIA integration of the WGS would also obligate a laboratory to implement a continuous performance measurement plan via an internal or external proficiency testing (PT) program. Such PT programs are under active development with the Global Microbial Identifier (GMI) network, the Genetic Testing Reference Materials Coordination Program (Get-RM), the Genome in a Bottle (GIAB) Consortium, and the CDC PulseNet NextGen being the most prominent [31, 49]. More generic standards have been proposed by the College of American Pathologists’ (CAP) molecular pathology checklist (MOL)[30]. The proposed quality standards include both live cultures as well as ‘sequence only’ formats for a comprehensive assessment of the WGS pipeline. Our validation set of isolates is amenable to both internal and external quality assurance testing. In preliminary internal PT, we were able to successfully assess the entire workflow and personnel performance (details not shown).

Microbial WGS remains a dynamic technology, and therefore, any validated pipeline is unlikely to remain static. For this reason, implementation of modular validation template becomes crucial for the seamless and timely introduction of changes to the ‘pipeline’, e.g. we had to carry-out several amendments to the protocol since its implementation in the laboratory. These included a new processing algorithm for highly-contagious pathogens and some adjustments to the data analysis algorithm. The changes were accomplished via minor modifications in the ‘pipeline’ with corroborative testing using developed by us modular validation template. We also performed a two-sequencer comparison to allow for processing of increased volume of samples (see the protocol for the correlation study in Supplementary Document 1).

The WGS report format continues to pose challenges. Reporting language was designed to be able to convey the WGS-based assay results to the end user with or without the extensive knowledge of WGS to avoid erroneous interpretation of the results by the final user and provide actionable data. Disclaimers are particularly important to guide the potential use of the data in clinical settings, e.g. a disclaimer that detection of antibiotic resistance genes by WGS do not guarantee resistance of the strain *in vivo* and that phenotypic susceptibility test is required to confirm antimicrobial resistance.

The study possesses certain limitations. Firstly, only a limited number of WGS-based assays were included into the validation study based on the most common PHL applications. Other types of WGS assays/analytics would have to be validated in a similar manner to determine the performance specifications, which are required to generate accurate and reproducible results, e.g. a threshold for the base calling accuracy of the platform, or a depth of coverage of specific genes. Secondly, not all validation set samples had available NCBI database entries to provide comparison sets. Thirdly, the absence of any eukaryotic pathogens in the current validation is another shortcoming and therefore, additional validation studies would be needed to implement a pipeline for the pathogenic fungi and parasites.

As the clinical and public microbiology community implements high-quality WGS, it would be opportune to consider the available models for the delivery of these services [54]. Since their inception, most WGS activities have taken place in the reference facilities with rather large supporting infrastructure. Although inevitable in the early stages, the centralization of services presents several challenges on the turnaround time and access to the specific expertise on the local population structure of a given pathogen, which are crucial for the management of infectious diseases at the local and regional levels. WGS services could now be delivered locally, more easily with the affordable sequencers, standardized reagents, and well-defined quality metrics. The local delivery model would also be more responsive to the needs of the target client and enhance the adoption of WGS across the healthcare systems. Another alternative is a hybrid model with complimentary central and local services to balance the need for speed with the advanced expertise and resources [54]. Two prominent examples of the hybrid models in the United States are the Food and Drug Administration (FDA) GenomeTrakr network for the tracking of food-borne pathogens, and the CDC Advanced Molecular Detection (AMD) initiative for the improved surveillance of infectious diseases [55, 56]. The AMD and GenomeTrakr frameworks rely on a participatory model with enhanced analysis, curation and data storage at a central site. However, these resource-intensive networks focus on few selected pathogens at present. Notably, there still remain significant challenges for the implementation of the comprehensive WGS services at the local level [48, 57]. It is hoped that the quality framework proposed in the present study would advance the localization of comprehensive WGS services in clinical and public health laboratories.

In summary, the salient achievements of this study included: 1) establishment of the performance specifications for WGS in the application to public health microbiology in accordance with CLIA guidelines for the LDTs, 2) the development of quality assurance (QA) and quality control (QC) measurements for WGS, 3) formatting of laboratory reports for end users with or without WGS expertise, 4) a set of pathogenic bacteria for further validations of WGS and multi-laboratory comparisons and, 5) development of an integrated workflow for the ‘wet bench’ and ‘dry bench’ parts of WGS.

## ACKNOWLEDGEMENT

This study was supported in part by the Epidemiology and Laboratory Capacity for Infectious Diseases Cooperative Agreement number 6 NU50CK000410-03 from the US Centers for Disease Control and Prevention.

The authors are thankful to Dr. Eija Trees, Chief of the PulseNet Next Generation Subtyping Methods Unit, at the Centers for Disease Control and Prevention for assistance in acquiring reference sequences for comparative study.

---

**Supplementary Figure 1.**
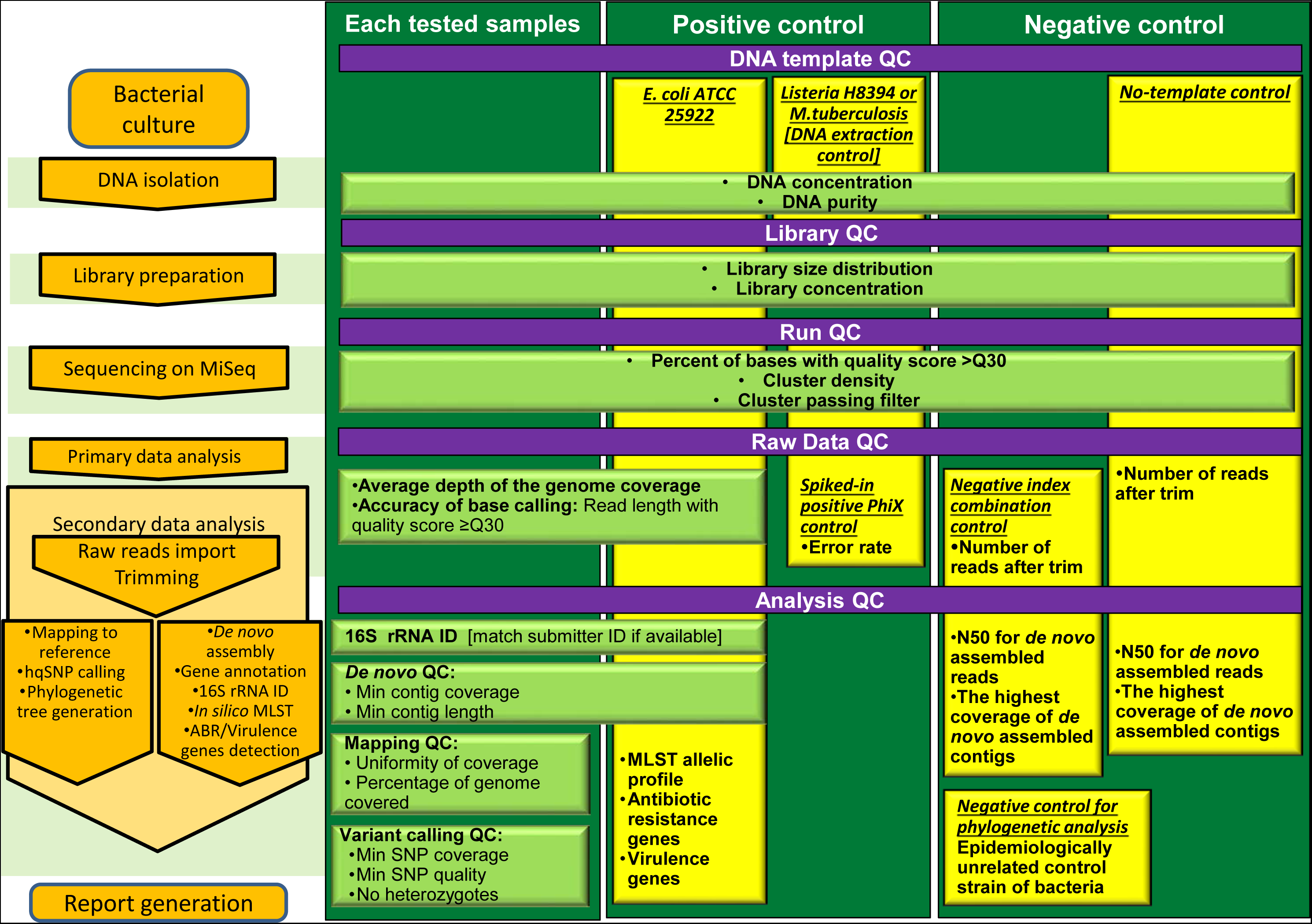
WGS quality control scheme.

## References

1. Didelot X, Bowden R, Wilson DJ, Peto TEA, Crook DW: Transforming clinical microbiology with bacterial genome sequencing. Nat Rev Genet 2012, 13(9):601–612.

2. Köser CU, Ellington MJ, Cartwright EJ, Gillespie SH, Brown NM, Farrington M, Holden MT, Dougan G, Bentley SD, Parkhill J: Routine use of microbial whole genome sequencing in diagnostic and public health microbiology. PLoS pathogens 2012, 8(8):e1002824.

3. Grad YH, Lipsitch M, Feldgarden M, Arachchi HM, Cerqueira GC, FitzGerald M, Godfrey P, Haas BJ, Murphy CI, Russ C et al: Genomic epidemiology of the Escherichia coli O104:H4 outbreaks in Europe, 2011. Proceedings of the National Academy of Sciences 2012, 109(8):3065–3070.

4. McGann P, Bunin JL, Snesrud E, Singh S, Maybank R, Ong AC, Kwak YI, Seronello S, Clifford RJ, Hinkle M et al: Real time application of whole genome sequencing for outbreak investigation – What is an achievable turnaround time? Diagnostic Microbiology and Infectious Disease 2016, 85(3):277–282.

5. Gordon NC, Price JR, Cole K, Everitt R, Morgan M, Finney J, Kearns AM, Pichon B, Young B, Wilson DJ et al: Prediction of Staphylococcus aureus Antimicrobial Resistance by Whole-Genome Sequencing. Journal of Clinical Microbiology 2014, 52(4):1182–1191.

6. Leopold SR , Goering RV, Witten A, Harmsen D, Mellmann A: Bacterial Whole-Genome Sequencing Revisited: Portable, Scalable, and Standardized Analysis for Typing and Detection of Virulence and Antibiotic Resistance Genes. Journal of Clinical Microbiology 2014, 52(7):2365–2370.

7. Goodwin S, McPherson JD, McCombie WR: Coming of age: ten years of next-generation sequencing technologies. Nature Reviews Genetics 2016, 17(6):333–351.

8. Roetzer A, Diel R, Kohl TA, Rückert C, Nübel U, Blom J, Wirth T, Jaenicke S, Schuback S, Rüsch-Gerdes S: Whole genome sequencing versus traditional genotyping for investigation of a Mycobacterium tuberculosis outbreak: a longitudinal molecular epidemiological study. PLoS Med 2013, 10(2):e1001387.

9. Gardy JL, Johnston JC, Sui SJH, Cook VJ, Shah L, Brodkin E, Rempel S, Moore R, Zhao Y, Holt R et al: Whole-Genome Sequencing and Social-Network Analysis of a Tuberculosis Outbreak. New England Journal of Medicine 2011, 364(8):730–739.

10. Eyre DW, Cule ML, Wilson DJ, Griffiths D, Vaughan A, O’Connor L, Ip CLC, Golubchik T, Batty EM, Finney JM et al: Diverse Sources of C. difficile Infection Identified on Whole-Genome Sequencing. New England Journal of Medicine 2013, 369(13):1195–1205.

11. Etienne KA, Roe CC, Smith RM, Vallabhaneni S, Duarte C, Escandón P, Castañeda E, Gómez BL, de Bedout C, López LF et al: Whole-Genome Sequencing to Determine Origin of Multinational Outbreak of Sarocladium kiliense Bloodstream Infections. Emerging Infectious Diseases 2016, 22(3):476–481.

12. Pallen MJ: Diagnostic metagenomics: potential applications to bacterial, viral and parasitic infections. Parasitology 2014, 141(Special Issue 14):1856–1862.

13. Onderdonk AB, Delaney ML, Fichorova RN: The Human Microbiome during Bacterial Vaginosis. Clinical Microbiology Reviews 2016, 29(2):223–238.

14. Pearce MM, Hilt EE, Rosenfeld AB, Zilliox MJ, Thomas-White K, Fok C, Kliethermes S, Schreckenberger PC, Brubaker L, Gai X et al: The Female Urinary Microbiome: a Comparison of Women with and without Urgency Urinary Incontinence. mBio 2014, 5(4).

15. Schubert AM, Rogers MAM, Ring C, Mogle J, Petrosino JP, Young VB, Aronoff DM, Schloss PD: Microbiome Data Distinguish Patients with Clostridium difficile Infection and Non-C. difficile-Associated Diarrhea from Healthy Controls. mBio 2014, 5(3).

16. Naccache SN, Federman S, Veeraraghavan N, Zaharia M, Lee D, Samayoa E, Bouquet J, Greninger AL, Luk K-C, Enge B et al: A cloud-compatible bioinformatics pipeline for ultrarapid pathogen identification from next-generation sequencing of clinical samples. Genome Research 2014, 24(7):1180–1192.

17. Hasman H, Saputra D, Sicheritz-Ponten T, Lund O, Svendsen CA, Frimodt-Møller N, Aarestrup FM: Rapid whole genome sequencing for the detection and characterization of microorganisms directly from clinical samples. Journal of Clinical Microbiology 2013.

18. Loman NJ, Constantinidou C, Christner M, et al.: A culture-independent sequence-based metagenomics approach to the investigation of an outbreak of shiga-toxigenic escherichia coli o104:h4. JAMA 2013, 309(14):1502–1510.

19. Endrullat C, Glökler J, Franke P, Frohme M: Standardization and quality management in next-generation sequencing. Applied & Translational Genomics 2016.

20. Salto-Tellez M, Gonzalez de Castro D: Next-generation sequencing: a change of paradigm in molecular diagnostic validation. J Pathol 2014, 234(1):5–10.

21. Goldberg B, Sichtig H, Geyer C, Ledeboer N, Weinstock GM: Making the Leap from Research Laboratory to Clinic: Challenges and Opportunities for Next-Generation Sequencing in Infectious Disease Diagnostics. mBio 2015, 6(6).

22. Kwong JC, McCallum N, Sintchenko V, Howden BP: Whole genome sequencing in clinical and public health microbiology. Pathology 2015, 47(3):199–210.

23. Luheshi LM, Raza S, Peacock SJ: Moving pathogen genomics out of the lab and into the clinic: what will it take? Genome Medicine 2015, 7(1):1–3.

24. Oliver GR, Hart SN, Klee EW: Bioinformatics for Clinical Next Generation Sequencing. Clinical Chemistry 2015, 61(1):124–135.

25. Fricke WF, Rasko DA: Bacterial genome sequencing in the clinic: bioinformatic challenges and solutions. Nat Rev Genet 2014, 15(1):49–55.

26. Wyres KL, Conway TC, Garg S, Queiroz C, Reumann M, Holt K, Rusu LI: WGS Analysis and Interpretation in Clinical and Public Health Microbiology Laboratories: What Are the Requirements and How Do Existing Tools Compare?

27. Rhoads DD, Sintchenko V, Rauch CA, Pantanowitz L: Clinical Microbiology Informatics. Clinical Microbiology Reviews 2014, 27(4):1025–1047.

28. Gargis AS, Kalman L, Lubin IM: Assuring the Quality of Next-Generation Sequencing in Clinical Microbiology and Public Health Laboratories. Journal of Clinical Microbiology 2016:JCM. 00949–00916.

29. Lefterova MI, Suarez CJ, Banaei N, Pinsky BA: Next-Generation Sequencing for Infectious Disease Diagnosis and Management: A Report of the Association for Molecular Pathology. The Journal of molecular diagnostics: JMD 2015, 17(6):623–634.

30. Aziz N, Zhao Q, Bry L, Driscoll DK, Funke B, Gibson JS, Grody WW, Hegde MR, Hoeltge GA, Leonard DGB et al: College of American Pathologists’ Laboratory Standards for Next-Generation Sequencing Clinical Tests. Archives of pathology & laboratory medicine 2014, 139(4):481–493.

31. Moran-Gilad J, Sintchenko V, Pedersen SK, Wolfgang WJ, Pettengill J, Strain E, Hendriksen RS: Proficiency testing for bacterial whole genome sequencing: an end-user survey of current capabilities, requirements and priorities. BMC Infectious Diseases 2015, 15(1):1–10.

32. Olson ND, Jackson SA, Lin NJ: Report from the Standards for Pathogen Identification via Next-Generation Sequencing (SPIN) Workshop. Standards in Genomic Sciences 2015, 10(119).

33. Gargis AS, Kalman L, Berry MW, Bick DP, Dimmock DP, Hambuch T, Lu F, Lyon E, Voelkerding KV, Zehnbauer BA: Assuring the quality of next-generation sequencing in clinical laboratory practice. Nature biotechnology 2012, 30(11):1033–1036.

34. Gargis AS, Kalman L, Bick DP, da Silva C, Dimmock DP, Funke BH, Gowrisankar S, Hegde MR, Kulkarni S, Mason CE et al: Good laboratory practice for clinical next-generation sequencing informatics pipelines. Nature biotechnology 2015, 33(7):689–693.

35. Duncavage EJ, Abel HJ, Merker JD, Bodner JB, Zhao Q, Voelkerding KV, Pfeifer JD: A Model Study of In Silico Proficiency Testing for Clinical Next-Generation Sequencing. Archives of pathology & laboratory medicine 2016, 140(10):1085–1091.

36. CLSI: Nucleic Acid Sequencing Methods in Diagnostic Laboratory Medicine: Approved Guideline-2d edition. MM09-A2. 2014.

37. Li H, Handsaker B, Wysoker A, Fennell T, Ruan J, Homer N, Marth G, Abecasis G, Durbin R, Genome Project Data Processing S: The Sequence Alignment/Map format and SAMtools. Bioinformatics 2009, 25(16):2078–2079.

38. Danecek P, Auton A, Abecasis G, Albers CA, Banks E, DePristo MA, Handsaker RE, Lunter G, Marth GT, Sherry ST et al: The variant call format and VCFtools. Bioinformatics 2011, 27(15):2156–2158.

39. Seemann T : Prokka: rapid prokaryotic genome annotation. Bioinformatics 2014, 30(14):2068–2069.

40. Cole JR, Wang Q, Fish JA, Chai B, McGarrell DM, Sun Y, Brown CT, Porras-Alfaro A, Kuske CR, Tiedje JM: Ribosomal Database Project: data and tools for high throughput rRNA analysis. Nucleic acids research 2014, 42(Database issue):D633–642.

41. Larsen MV, Cosentino S, Rasmussen S, Friis C, Hasman H, Marvig RL, Jelsbak L, Sicheritz-Ponten T, Ussery DW, Aarestrup FM et al: Multilocus sequence typing of total-genome-sequenced bacteria. J Clin Microbiol 2012, 50(4):1355–1361.

42. Joensen KG, Scheutz F, Lund O, Hasman H, Kaas RS, Nielsen EM, Aarestrup FM: Real-time whole-genome sequencing for routine typing, surveillance, and outbreak detection of verotoxigenic Escherichia coli. J Clin Microbiol 2014, 52(5):1501–1510.

43. CLSI: Performance Standards for Antimicrobial Susceptibility Testing; Twenty-Fifth Informational Supplement. CLSI document M100-S25. Wayne, PA: CLinical and Laboratory STandards Institute; 2015. 2015.

44. Harris SR, Cartwright EJ, Torok ME, Holden MT, Brown NM, Ogilvy-Stuart AL, Ellington MJ, Quail MA, Bentley SD, Parkhill J et al: Whole-genome sequencing for analysis of an outbreak of meticillin-resistant Staphylococcus aureus: a descriptive study. The Lancet Infectious diseases 2013, 13(2):130–136.

45. Leekitcharoenphon P, Nielsen EM, Kaas RS, Lund O, Aarestrup FM: Evaluation of whole genome sequencing for outbreak detection of Salmonella enterica. PloS one 2014, 9(2):e87991.

46. CLSI: Molecular Methods for Bacterial Strain Typing; Approved Guideline, MM11-A. 2007.

47. Richards S , Aziz N, Bale S, Bick D, Das S, Gastier-Foster J, Grody WW, Hegde M, Lyon E, Spector E et al: Standards and guidelines for the interpretation of sequence variants: a joint consensus recommendation of the American College of Medical Genetics and Genomics and the Association for Molecular Pathology. Genet Med 2015, 17(5):405–423.

48. Lesho E, Clifford R, Onmus-Leone F, Appalla L, Snesrud E, Kwak Y, Ong A, Maybank R, Waterman P, Rohrbeck P et al: The Challenges of Implementing Next Generation Sequencing Across a Large Healthcare System, and the Molecular Epidemiology and Antibiotic Susceptibilities of Carbapenemase-Producing Bacteria in the Healthcare System of the U.S. Department of Defense. PloS one 2016, 11(5):e0155770.

49. Kalman LV, Datta V, Williams M, Zook JM, Salit ML, Han J-Y: Development and Characterization of Reference Materials for Genetic Testing: Focus on Public Partnerships. Annals of Laboratory Medicine 2016, 36(6):513–520.

50. Lindsey RL, Pouseele H, Chen JC, Strockbine NA, Carleton HA: Implementation of Whole Genome Sequencing (WGS) for Identification and Characterization of Shiga Toxin-Producing Escherichia coli (STEC) in the United States. Frontiers in Microbiology 2016, 7(766).

51. Pankhurst LJ, del Ojo Elias C, Votintseva AA, Walker TM, Cole K, Davies J, Fermont JM, Gascoyne-Binzi DM, Kohl TA, Kong C et al: Rapid, comprehensive, and affordable mycobacterial diagnosis with whole-genome sequencing: a prospective study. The Lancet Respiratory Medicine 2016, 4(1):49–58.

52. Walker TM, Kohl TA, Omar SV, Hedge J, Del Ojo Elias C, Bradley P, Iqbal Z, Feuerriegel S, Niehaus KE, Wilson DJ et al: Whole-genome sequencing for prediction of Mycobacterium tuberculosis drug susceptibility and resistance: a retrospective cohort study. The Lancet Infectious Diseases 2015, 15(10):1193–1202.

53. Zankari E, Hasman H, Cosentino S, Vestergaard M, Rasmussen S, Lund O, Aarestrup FM, Larsen MV: Identification of acquired antimicrobial resistance genes. The Journal of antimicrobial chemotherapy 2012, 67(11):2640–2644.

54. Arnold C : Considerations in centralizing whole genome sequencing for microbiology in a public health setting. Expert Review of Molecular Diagnostics 2016, 16(6):619–621.

55. Auffray C, Caulfield T, Griffin JL, Khoury MJ, Lupski JR, Schwab M: From genomic medicine to precision medicine: highlights of 2015. Genome Medicine 2016, 8(1):12.

56. Allard MW, Strain E, Melka D, Bunning K, Musser SM, Brown EW, Timme R: Practical Value of Food Pathogen Traceability through Building a Whole-Genome Sequencing Network and Database. Journal of Clinical Microbiology 2016, 54(8):1975–1983.

57. Robilotti E, Kamboj M: Integration of Whole-Genome Sequencing into Infection Control Practices: the Potential and the Hurdles. Journal of Clinical Microbiology 2015, 53(4):1054–1055.

